# Transposable Elements are Dysregulated in Brains of Individuals with Major Depressive Disorder

**DOI:** 10.1101/2025.01.22.634143

**Authors:** Natalie L. Truby, Corinne Smith, Peter J. Hamilton

## Abstract

Transposable elements (TEs) are repetitive DNA sequences capable of being transcribed and reintegrated, or transposed, into distinct loci throughout the genome. While thought to be largely transcriptionally silenced in brain, TE transcription is increasingly recognized as dynamic and involved in human health and disease states, including in disorders of the brain. In this study, we annotated TE transcripts in publicly available RNA sequencing (RNAseq) of postmortem human brain tissue to investigate the expression profile of TE transcripts in individuals with Major Depressive Disorder (MDD) compared to healthy controls. Our findings reveal a robust and uniform downregulation of TE transcript expression in the brains of subjects with MDD relative to controls, this occurs most prominently in the orbitofrontal cortex (OFC) brain region, and MDD differentially impacts this TE expression by age and sex. This work points to the aberrant transcription of cortical TEs as a potentially overlooked molecular signature of MDD.

## Introduction

A growing body of evidence implicates the transcriptional control of brain transposable elements (TEs) in neuropsychiatric illness and related complex behaviors^1–6^. Originally considered “junk DNA,” TEs are repetitive sequences that have the ability to self-replicate and transpose within the host genome^7^. Recent studies have revealed a dynamic interplay between TEs and various cellular processes implicated in brain diseases, including gene expression, genome stability and structure, and epigenetic status^4,6–10^. Further, several lines of evidence suggest that KRAB-zinc finger protein (KZFP) transcription factors (TFs) are a major molecular mechanism dedicated to binding TEs and repressing TE transcription^11–13^, which has enabled domestication of TEs to form *cis*-regulatory genomic elements to facilitate more complex gene expression patterns^11–17^.

Our group has previously studied an individual KZFP whose expression in mouse prefrontal cortex (PFC) is activated by chronic stress and is protective against stress-induced social deficits^18^. By manipulating the PFC functions of this KZFP, we further established a relationship between this KZFP, cortical TE transcriptional control, and the social cognition necessary for complex pro-social behavior^19^. Given that this prior work involved synthetic molecular manipulations in laboratory rodents, we wondered if brain TE stability is capable of being naturally altered in human disease states. Here, we interrogated TE expression in the brains of individuals with Major Depressive Disorder (MDD). We elected to study MDD since chronic stress is a risk factor for developing MDD^20^ and social deficits are a common symptom of the disorder^21^. While genetic and environmental factors have been implicated in the etiology of MDD^20,22^, the exact molecular mechanisms that produce MDD symptoms remain elusive, and current pharmacotherapies are suboptimal. Mechanistic insights into TE biology in brain disease states have grossly ignored due to the fact that standard RNAseq analytical pipelines routinely exclude reads with multiple alignments in the reference genome. As a consequence of transposition, TE transcript reads often align to multiple TE-related genomic loci, and for this reason and TE transcript expression is discarded in nearly all RNAseq studies. Here, we employed previously developed and validated bioinformatic tools to specifically capture and characterize brain TEs in published RNAseq data performed on multiple brain regions of people with MDD and matched controls^23^.

## Results

We observe that TE transcripts are uniformly decreased in the brains of individuals with MDD. Our results demonstrated that every one of the 566 detected significant DETEs (adjusted p-value ≤ 0.05) was downregulated in the brains of MDD patients when compared to their healthy counterparts (Fig. 1A, B). Specifically, we observed that all three classes of TEs were affected, with the long terminal repeat (LTR) transposon family constituting the largest fraction of the downregulated TEs (50.2%) (Fig. 1C). Given that non-LTR transposons are most ubiquitous in the mammalian genome^24^ and thought to be the only human TE class still capable of transposition^25^, it was unexpected that non-LTR class of TEs was least represented in the DETE list. These data indicate that the transcripts corresponding to LTR and DNA TE classes, which are thought to be largely transcriptionally inactive, are disproportionately affected in MDD brains.

**FIGURE 1.**
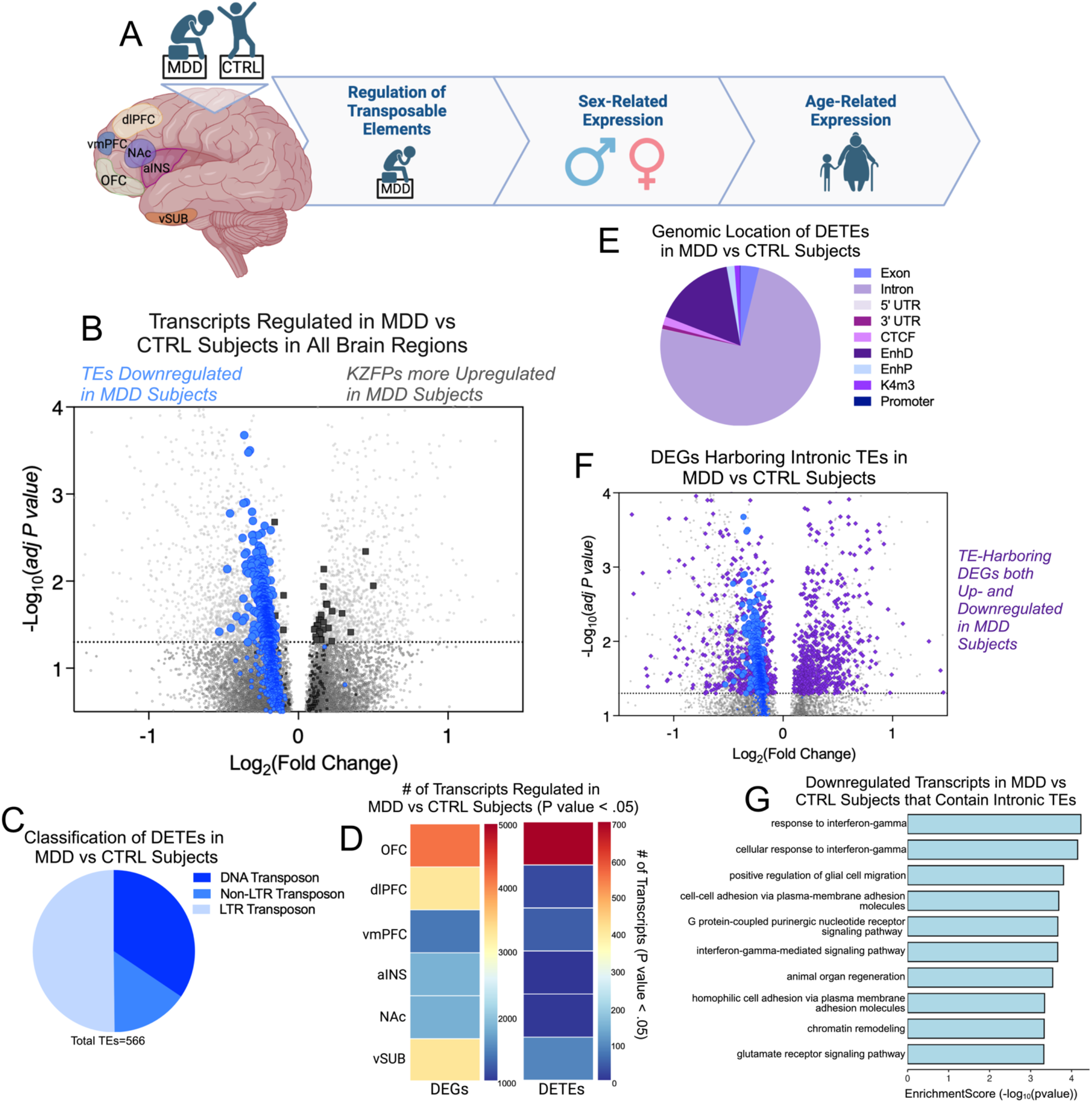
Transposable element transcripts are uniformly downregulated in the brains of MDD patients. **(A)** Schematic representing experimental design: RNAseq data^23^ from six different postmortem brain regions from patients with MDD and healthy matched controls is used to annotate regulation of transposable elements (TEs), sex-related TE expression and age-related TE expression. Cartoon made in Biorender. **(B)** Volcano plot depicting expression profiles of differentially expressed genes (DEGs) and differentially expressed TEs (DETEs) in MDD vs CTRL brains. Analysis of publicly available RNAseq^23^ of human postmortem brain tissue from six brain regions combined: orbitofrontal cortex (OFC; BA11), dorsolateral PFC (dlPFC; BA8/9), ventromedial PFC (vmPFC; BA 25), anterior insula (aiNS), nucleus accumbens (NAc) and ventral subiculum (vSUB), using a model that considers all covariates (sex, age, medication, smoking), reveals that many significant DETEs (blue circles) are downregulated, and none are upregulated in MDD vs CTRL. Significantly DEGs encoding KZFPs (dark grey squares) are primarily upregulated in MDD vs CTRL. Canonical DEGs are represented by light grey circles. Significance cutoff set to adjusted *p* value < .05. n = 26 MDD postmortem samples per brain region (13 males; 13 females), n = 22 CTRL postmortem samples per brain region (13 males; 9 females) **(C)** Representation of TE families for MDD-specific DETEs. Most (50.2%) of the observed MDD-specific DETEs belong to the LTR transposon family, 15.4% belong to the non-LTR transposons family, and 34.45% belong to the DNA transposon family. **(D**) Heat map showing number of significant (p value < .05) DEGs and DETE transcripts in each brain region **(E)** A majority of the observed MDD-specific DETEs (74.5%) map to TE sequences localized to intronic regions of the genome. **(F)** Volcano plot depicting expression profiles of DETEs (blue circles) and TE-harboring DEGs (purple diamonds). TE-harboring DEGs are up- and downregulated, while TE transcripts are only downregulated. Significance cutoff set to adjusted *p* value < .05. **(G)** GO analysis reveals enrichment of immune and interferon signaling genes in down-regulated TE harboring genes.

Given the established repressive action of KZFP TFs at TEs, we also examined the expression patterns of KZFP encoding genes within our dataset. Interestingly, we found that KZFP transcript levels skewed towards up-regulated in the brains of individuals with MDD compared to healthy controls (Fig. 1B), which might hint at a mechanistic relationship between up-regulation of KZFP TFs and the hyper-repression of TEs that is observed in MDD brain.

To determine the MDD-associated TE expression pattern within individual brain regions, we conducted individual analyses of DETEs and DEGs within the six studied brain regions: the orbitofrontal cortex (OFC; BA11), dorsolateral prefrontal cortex (dlPFC; BA8/9), ventromedial prefrontal cortex (vmPFC; BA25), anterior insula (aINS), nucleus accumbens (NAc), and ventral subiculum (vSUB). The OFC exhibited the most pronounced impact of DETEs (all downregulated) as well as the highest number of DEGs (Fig. 1D), while all other brain regions possessed fewer DETEs.

To elucidate the potential genomic origin of the observed DETEs, in our human reference genome we annotated genomic features that corresponded to regions of aligned to TE reads (Fig. 1E). A majority (74.5%) of the MDD-specific DETEs could be mapped to TE sequences located in gene intron regions. Introns are a noted hotspot for harboring TE sequences, and intronic TEs can alter gene splicing patterns, expression levels, and function^26–28^.

To explore the possibility that intronic TE dysregulation affects the expression of the gene in which the intronic TE resides, we next identified if any TE-harboring DEGs were significantly affected in these datasets (Fig. 1F). We identified the gene IDs known to contain intronic TE sequences of affected DETEs and queried if the associated DEG was also significantly affected in MDD brain.

These TE-harboring DEGs were both up and downregulated in MDD brain (Fig. 1G), pointing to the possibility that not all TE-harboring genes experience TE dysregulation or that the consequence of this intronic TE dysregulation does not uniformly impact the expression of the proximal gene. This points to a mechanism where the transcriptional control of intronic TEs is at least partially independent from the transcription of the gene in which it resides. As the significant DETEs in the brain of MDD subjects are uniformly downregulated, we chose to analyze the downregulated DEGs harboring intronic TEs with the thought that these could be the most sensitive genes to TE dysregulation. In conducting a gene ontology (GO) analysis of these TE-harboring DEGs, we see enrichment of biological pathways related to immune function, particularly interferon gamma signaling (Fig. 1H). These findings further contribute to the growing body of literature linking immune function (particularly interferon signaling) with social behaviors and neuropsychiatric disorders^29–31^, including prior work from our group^19^. Further studies are required to establish any direct mechanistic relationship between transcriptional control of intronic TEs and its effect on proximal gene expression.

We next investigated how sex and age influence the expression of DETEs in the brains of individuals with MDD compared to healthy matched controls. Our analyses revealed that TE expression was consistently higher in males than in females within both the MDD and control groups (Fig. 2A-C). Notably, MDD pathology appears to enhance existing sex-related differences in TE expression, as the MDD group has 68 significant upregulated DETEs in males vs. females, while the control group has 19. Moreover, we noted that 84.2% of the sex-specific DETEs identified in the control group were also present in the MDD group (Fig. 2F), and the MDD group exhibited a predominance of unique DETEs that were not observed in controls (Fig. 2F). This suggests that MDD not only maintains the established sex-related differences in TE expression but also enhances them, resulting in a distinct TE expression profile in affected individuals, and potentially contributing to the sex-specific prevalence and symptomatology observed in MDD^32^. It is well appreciated that TE regulation is increased in aging and in the brains of older individuals^8,28,33^. However, we sought to observe if age-induced changes in TE expression vary in the brains of those with MDD. Analysis of TE expression across different age groups (age ≤ 46 (younger) vs. age > 46 (older)) revealed that TEs were downregulated in younger subjects in the control group, as expected (Fig. 2D). However, in the MDD group, there was no significant age-related pattern in TE expression (Fig. 2E).

**FIGURE 2.**
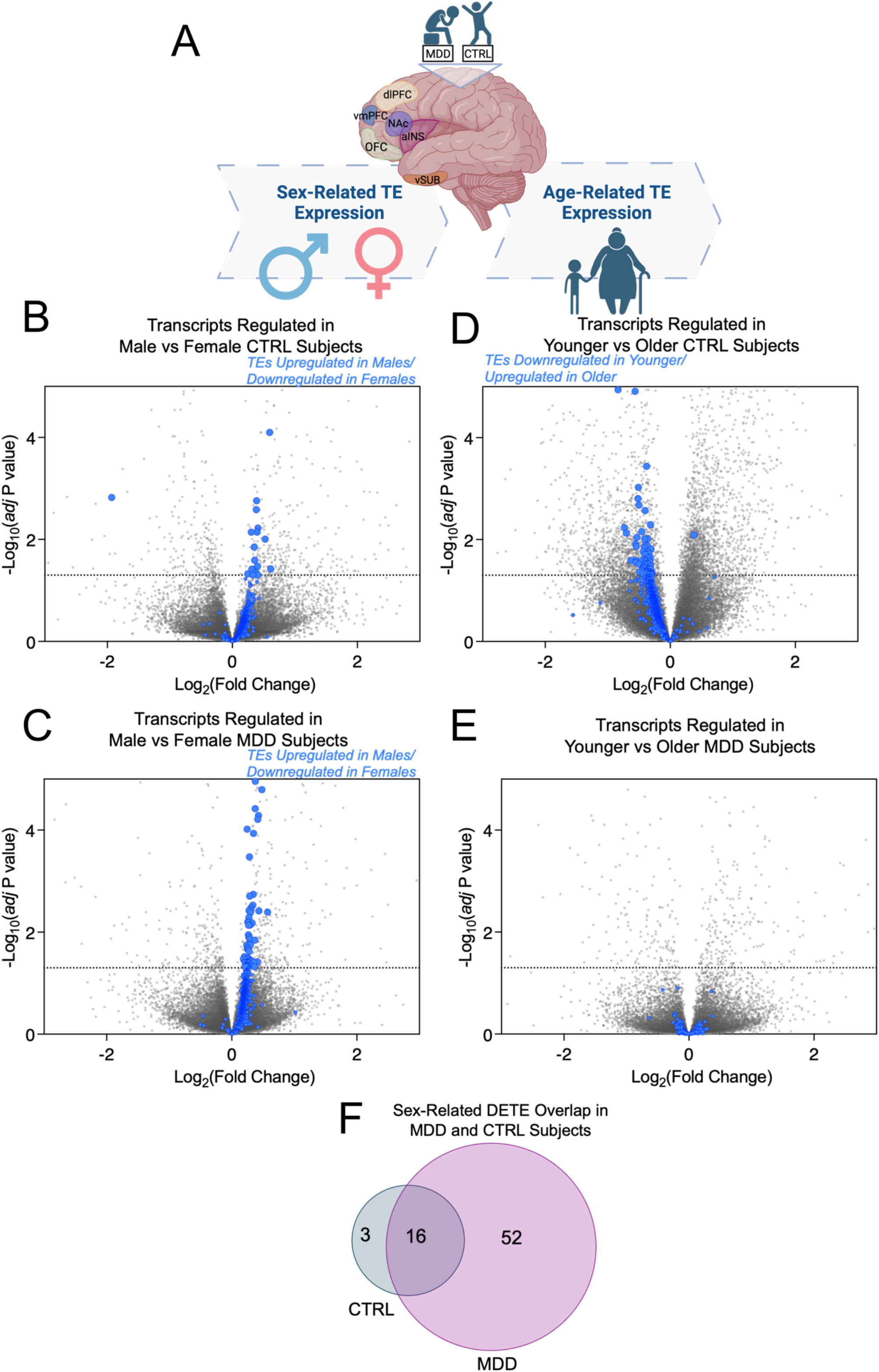
MDD alters age and sex differences in brain TE expression. **(A)** Schematic representing experimental design. Cartoon made in Biorender. **(B)** Volcano plot depicting expression patterns of DETEs in males vs. females in CTRL group. 18 significant DETEs (blue circles) are upregulated and only one is downregulated. Canonical DEGs are represented by light grey circles. Significance cutoff set to adjusted *p* value < .05. n = 13 males, n = 9 females. **(C)** Volcano plot depicting expression patterns of DETEs in males vs. females in MDD group. 68 significant DETEs (blue circles) are uniformly upregulated in males vs. females within the MDD group. Canonical DEGs are represented by light grey circles. Significance cutoff set to adjusted *p* value < .05. n = 13 males, n = 13 females **(D)** Volcano plot depicting age-specific expression patterns of DETEs in individuals with age ≤ 46 (younger) vs. age > 46 (older) in CTRL group. Median age in CTRL group = 46. 78 significant DETEs (blue circles) are downregulated and only one is upregulated. Canonical DEGs are represented by light grey circles. Significance cutoff set to adjusted *p* value < .05. n = 12 age ≤ 46, n = 10 age > 46. **(E)** Volcano plot depicting age-specific expression patterns of DETEs in individuals with age ≤ 47 (younger) vs. age > 47 (older) in MDD group. Median age in MDD group = 47. DETE (blue circles) expression is not impacted by age in MDD group. This indicates that there is age-related DETE expression within the MDD group but not CTRL group. Canonical DEGs are represented by light grey circles. Significance cutoff set to adjusted *p* value < .05. n = 13 age ≤ 47, n = 13 age > 47. **(F)** Venn diagram displaying sex-specific DETE overlap between MDD and CTRL groups. MDD augments sex-specific DETE expression. Significance cutoff set to adjusted *p* value < .05.

## Discussion

Here, we present novel insights into the role of brain TE transcriptional regulation in MDD. We reveal a uniform downregulation of TEs in the brains of affected individuals compared to healthy controls.

This finding points to the transcriptional control of brain TEs as dynamic and a potentially overlooked molecular signature of neuropsychiatric disorders. TEs are thought to be transcriptionally silent in most tissues, with their expression tightly regulated to prevent genome instability^28,34–36^. In MDD, the downregulation of TE transcripts across multiple brain regions may reflect a broader dysregulation of genomic stability and transcriptional control.

The observation that OFC experienced the most significant downregulation of TEs supports the notion that some specific brain regions are more susceptible to transcriptional dysregulation in MDD^37–39^. The OFC has undergone significant expansion and reorganization in mammals, particularly in primates, likely supporting more sophisticated cognitive processes^40^. Similarly, TEs have played a crucial role in shaping genome structure and diversity, contributing to the evolution of species-specific traits^41,42^, such as MDD. The interplay between these two evolutionary processes is notable, as TE activity within regulatory regions of the genome may have facilitated the adaptive changes in brain structures like the OFC^35,43^.

The observation of higher TE expression in males compared to females in both the MDD and control groups is noteworthy. Sex differences in TE expression are not well understood, but emerging evidence suggests that sex hormones may play a regulatory role in TE activity^44–47^, potentially explaining why males and females exhibit different TE expression profiles. In MDD, these sex differences are potentiated, with the MDD group displaying an even greater male bias in TE expression than the control group. This enhanced male bias in MDD may contribute to our understanding of the higher prevalence and different symptomatology of MDD in females^23,32,48^.

As individuals age, the genome becomes more prone to instability, and TE mobilization increases, leading to genetic changes that may impact brain function^8,28,33^. In the control group, we observed a typical downregulation of TE expression in younger individuals, which may reflect age-dependent changes in TE activity (Fig. 2D). However, in the MDD group, this age-related pattern was absent (Fig. 2E), suggesting that the brain in MDD may exhibit characteristics of an “aged” brain, in terms of TE expression. This is consistent with the idea that MDD could accelerate certain age-related molecular processes^49,50^, such as the dysregulation of TEs, and may contribute to the cognitive and neurobiological deficits observed in older adults with depression^51,52^.

TE-rich regions of the genome are potent sources of *cis-*regulatory elements^53^. In healthy brains, TE stability is appreciated as a modulator of gene expression^28^, providing the necessary transcriptional flexibility for adaptive responses to environmental challenges. In contrast, the dysregulation of brain TEs seen in MDD brain may impair key transcriptional flexibility and could contribute to the vulnerability and pathogenesis of MDD symptomology^54^. Our work in animal models indicate that directly dysregulating cortical TEs with a synthetic KZFP impedes interferon and immune gene expression and produces social deficits^18^. This suggests that the dysregulation of brain TEs seen here in MDD may actually be causative in the pathogenesis of the disorder, especially as it relates to the emergence of MDD-associated social deficits. Overall, our findings reveal a previously overlooked dynamic transcriptional control of brain TEs in people with neuropsychiatric disorders and may offer novel insights into the etiology and treatment of MDD.

## Materials and Methods

Raw RNAseq datasets were downloaded from NCBI GEO GSE102556. Details on methods and analyses are available as SI Appendix.

## Acknowledgements

This work was supported by the NIH R00DA045795, P30DA033934, R34DA061267, R01DA058958, R01DA058089, and Blick Scholar funds to PJH, as well as F31MH133309 to NLT.

## Supplemental Information (SI) Appendix

### Extended Materials and Methods

We conducted a comprehensive analysis of publicly available RNAseq data from human postmortem brain tissue^23^ across six brain regions: the orbitofrontal cortex (OFC; BA11), dorsolateral prefrontal cortex (dlPFC; BA8/9), ventromedial prefrontal cortex (vmPFC; BA25), anterior insula (aINS), nucleus accumbens (NAc), and ventral subiculum (vSUB). Our annotation of TEs was performed using TETranscripts^55^ allowing us to generate a list of differentially expressed TEs (DETEs), as well as differentially expressed genes (DEGs), in individuals diagnosed with MDD compared to matched healthy controls (CTRLs). The analysis included a total of 26 postmortem samples from MDD subjects (comprising 13 males and 13 females) and 22 control samples (13 males; 9 females), and used a model that considers all covariates (including sex, age, medication, smoking).

### RNA-seq data preprocessing and quality control

Publicly available RNAseq datasets were downloaded from NCBI GEO GSE102556. Raw RNA-seq FASTQ files were subjected to quality control using FastQC (version 0.11.9) to assess read quality. Adapter sequences were removed using Trimmomatic (version 0.39) with further trimming off low-quality bases (10 bases from the head, and 3 base from the tail, respectively). In addition, we used a sliding window to trim off the bases with average base quality lower than 20. Reads shorter than 16 (half of 50-10-3) were also be dropped.

### RNAseq read alignment and TE/gene quantification

After the quality control, high-quality reads were aligned to the Homo sapiens GRCh38 reference genome for human data available on Ensembl using STAR (version 2.7.11a) with the recommended parameters. We employed TEtranscripts (version 1.09)^55^ in the quantification of both gene and TE expression levels by integrating genomic and RepeatMasker annotations (Hammell lab).

### Differential Expression Analysis

The RNAseq count data were normalized, and dispersion estimates were obtained according to DESeq2’s standard pipeline. We employed a model in which age, sex, medication and smoking history were adjusted as covariates. Medication type was categorized into Alzheimer’s disease, anti-convulsant, antipsychotic, lithium, and no medication. Smoking history was categorized into heavy, moderate, and non-smoker. When comparing MDD vs CTRL samples, and in looking at sex and age, our DE analysis model utilized the whole-brain counts with all brain region samples combined. The RNAseq counts from different brain regions of the same subject were averaged to make the (pseudo-) whole-brain counts. Wald tests were performed to determine the significant levels of differentially expressed genes (DEGs) or TEs (DETEs) between the two clinical conditions (MDD vs. CTRL). The DEGs and DETEs were considered significant if the absolute log2 fold change were greater than 1 and the false discovery rate (FDR) less than 0.05. Volcano plots with these DEGs and DETEs were assembled in GraphPad Prism 10.

### Genomic annotation of TEs and potential regulated genes

We annotated the TEs of overlapping genomic features. The genomic features included in this analysis were the promoter regions, enhancers, histone H3K4 tri-methylation sites and CTCF binding sites revealed by ENCODE project, as well as the exon, intron, UTR from GRCh38 gene annotations. The overlapping genomic features were identified based on a minimum of 1 bp overlap of the chromosomal position between the observed TEs and the genomic features. The genes with introns overlapping with TEs were identified directly from gene annotations. The genes that are potentially regulated by the overlapping enhancers were revealed by analyzing the data of various high-throughput experiments, e.g. histone modifications, DNase-seq, or ChIA-PET. Downregulated DEGs in MDD vs CTRL conditions (p < 0.05 cutoff) that harbor intronic TEs were analyzed using SRplot gene ontology generator^56^.

